# Generalist flowers recover rapidly in grassland restoration: The importance of floral traits in community reassembly

**DOI:** 10.1101/2025.06.24.661246

**Authors:** Gaku S. Hirayama, Hiroshi S. Ishii, Taisei W. Ueno, Kenta Tanaka, Atushi Ushimaru

## Abstract

Community assembly process, through biotic and abiotic filtering of species traits, is crucial for understanding the recovery of biodiversity and ecosystem function in restored ecosystems. While previous studies have highlighted very slow recovery of pollination networks in ecological restoration, the reassembly process of plant communities focusing on floral traits that influence pollination networks remains underexplored. We hypothesised that plant species with flowers that allow visits by various pollinators (generalist flowers) would recover more easily than those with flowers which limit pollinators (specialist flowers) during grassland restoration. We investigated floral trait assemblages, interaction patterns with pollinators, and pollination success for each floral trait (floral tube and colour) across 28 pollination networks. We examined their temporal change patterns within restored grasslands (2-75 years old) and compared them with reference old grasslands (>165 years old). Our results revealed that generalist (open/shallow and white/yellow) flowers have recovered rapidly, interacting with many pollinator species with similar traits in the early stages of restoration. In contrast, specialist (long-tubed and blue/purple) flowers exhibited more delayed recovery due to low pollination success, taking approximately 75 years for the recovery to the levels of reference old grasslands. Importantly, we found that functional specialisation was crucial for the recovery of these specialist flowers. Corresponding to temporal changes in floral trait assemblages, pollinator composition has shifted from dipteran dominance to a greater abundance of long-tongued bees, with interaction patterns of each trait group approaching those of old grasslands. Our findings underscore the pivotal role of floral trait reassembly in influencing temporal changes in pollinator community, network structure, and pollination function during grassland restoration. By integrating trait-based framework, this study provides novel insights into the co-reassembly process of plants and pollinators in restored ecosystems.

## INTRODUCTION

Trait-based community assembly, which focuses on how local communities establish and change through species dispersal, abiotic, and biotic filtering based on species traits, is a longstanding theme in ecology. Recently, this trait-based framework has been applied in community restoration to understand which traits are more likely to be important for species recovery in human-managed ecosystems (Funk, 2021; Balazs et al., 2021). Trait-based reassembly process is particularly crucial for ecosystems with slow natural recovery rates, where understanding which traits promote successful establishment and recovery of original flora and fauna can guide anthropogenic interventions to accelerate community restoration.

In terrestrial ecosystems, grasslands have been a primary focus of ecological restoration (Dengler et al., 2014). Natural and semi-natural grasslands have been rapidly lost and degraded worldwide (Dengler et al,. 2014), despite their high biodiversity and essential ecosystem function, such as pollination, carbon cycling, and water resource supply (Zhao et al., 2020; Hirayama et al., 2025). Recent works on grassland restoration have mainly focused on how seed dispersal modes and key environmental factors (soil physicochemical properties and light availability) affect plant community reassembly after restoration onset (Baer et al., 2019; Yaida et al., 2024a,b). Meanwhile, although mutualistic biotic interactions with other trophic levels, such as plant-pollinator and plant-microbe interactions, have significant effects on plant reproduction and growth (Cariveau et al., 2020; Abrahao et al., 2022), they have been largely ignored in plant community reassembly of restored grasslands (Hirayama et al., 2025). Furthermore, the recovery of mutualistic interactions may lead to the restoration of ecosystem function. Thus, trait-based community reassembly must be updated to explicitly consider the functional role of traits in recovering these mutualistic trophic interactions in grassland restoration.

Pollination interactions are key trophic interactions for community reassembly in restored grasslands (Kaiser-Bunbury et al., 2017; Hirayama et al., 2025), because many flowering forbs depend on wild pollinators for their reproduction in temperate regions (Ollerton et al., 2011). Insect-pollinated plant species usually take longer time to recover after grassland restoration with seeding, compared to wind-pollinated plants (Albert et al., 2021). In fact, restored grasslands exhibit low pollination function even after several decades of restoration due to altered pollinator compositions and generalised pollination networks, which may lead to restoration delay (Hirayama et al., 2025). Thus, the recovery of pollinator community and their interactions with plants is critical in plant community reassembly. As primary key traits shaping plant-pollinator interactions, floral shape and colour may function as important drivers for community assembly in grassland ecosystems (Ishii et al., 2025). While a few studies have highlighted the role of pollination in ecological restoration (Kaiser-Bunbury et al., 2017; Hirayama et al., 2025), the reassembly processes of these trait composition in restored plant-pollinator interactions remain completely unexplored.

Here, we hypothesise that floral traits attracting a broader range of pollinators are more advantageous for establishing success of given plants compared to those adapted to specific pollinators in the early stages of grassland restoration, given the potentially limited or unpredictable pollinator communities in recently restored habitats. For instance, corolla tubes (or spurs) significantly limit pollinator accessibility: deep flowers are usually pollinated by specialist animals with long proboscises, while open or shallow flowers are accessible to a wider variety of pollinators (Stang et al., 2009; Lazaro et al., 2020; Hiraiwa and Ushimaru, 2017). Likewise, flower colours can influence pollinator attraction patterns; white and yellow colours, which reflect a wider range of wavelengths, are generally visible to and visited by a broad range of pollinators. In contrast, blue and red colours are perceived and preferred by specific pollinator groups due to their narrower reflectance range (Chittka et al., 2001; Chen et al., 2020; Ishii et al., 2025). Such generalist flowers (open/shallow and white/yellow flowers), despite facing risks of heterospecific pollen deposition, may receive higher overall pollination success in young restored grasslands where pollinator communities are still developing and limited in diversity and composition. Consequently, plant species with generalist floral traits are likely to recover more quickly in young restored grasslands while recovery of those preferred by specialist pollinators may take longer during the process of plant community reassembly.

In this study, we examine the reassembly pattern of community-wide floral traits and whether the degree of specialisation increases with changes in floral trait composition during grassland restoration on ski slopes. In our study system, some ski grasslands were reestablished after forest clear-cutting following temporal forestation, and we consider them as examples of restored grasslands while others were converted from old pastures and serve as reference old grasslands (Yaida et al., 2024a, b; Hirayama et al., 2025). We examined 28 pollination communities in 7 restored grasslands with different reconstruction times (2-75 years ago) and 7 reference old grasslands (at least 165 years old). Our previous study revealed that younger restored grasslands with low insect-pollinated plant richness have a high degree of network-level generalisation and that restored grasslands become more specialised as plant richness increases over several decades (Hirayama et al., 2025). In the previous study, however, we did not examine trait-based reassembly of plant-pollinator interactions in the restored grasslands. Here, we test our hypothesis that plant species with generalist-attracting floral traits recover more quickly in younger restored grasslands compared to those adapted to specialist pollinators by analysing chronosequence changes in community-wide floral tube (spur) length and colour composition. We further investigate changes in pollinator composition and pollination success for each floral trait group over the years following restoration initiation.

## MATERIALS & METHODS

The study area, study plots, and survey methods of plants, pollinators, and their interactions are described in detail in Hirayama et al., (2025). Therefore, in the following, only a brief overview is provided.

### Study area, ski slopes, and plots

Flower-pollinator surveys were performed in 2021 and 2022 on 14 ski slopes at three ski resorts in the Sugadaira Highland in Nagano Prefecture, center of Japan, five slopes at Davos, seven slopes at Taro, and two slopes at Omatsu (36°51’–54’N, 138°31’–36’E, approximately 1,300–1,500 m a.s.l.). The climate and geographical information for study sites is presented in a previous study (Hirayama et al., 2025). All study grassland has been managed by annual mowing in autumn (Yaida et al., 2024a, b). For restored grasslands, years since grassland restoration onset (grassland age) was estimated using past aerial photographs (Inoue et al., 2021; Yaida et al., 2024a).

### Flower-pollinator networks

We established a 1000 m² (5 × 200 m) belt plot for each grassland site. Five seasonal surveys were conducted in 14 sites, approximately once a month, on sunny and warm days from May to September in 2021 and 2022. Mowing timing differed slightly among the grasslands, from late August to early September; therefore, two sites and one site were surveyed four times a year in 2021 and 2022, respectively. During each seasonal survey, we recorded all plant species with insect-pollinated flowers and counted the number of flowers for each species in each plot. We then recorded flower visitors who touched the stigmas and anthers as ‘pollinators’. Each survey comprised four observations (two in the morning, 0800–1200 h and two in the afternoon, 1200–1600 h). Specimens were kept in our laboratory and as far as possible identified to the species level. In total, we constructed 28 quantitative plant-pollinator networks (14 sites × 2 years). We summarized the annual data collected from each site as a unit to calculate the community and generality parameters for subsequent analyses.

To estimate the flower abundance of each species at each site and during each survey, we measured several flower parameters to calculate the visual (attraction) size (mm²) of the three sampled flowers for each plant species, following Hiraiwa and Ushimaru (2017). For each site and survey, flower abundance of each species was calculated as the total visual size of the flowers (mean visual size multiplied by the number of flowers). Furthermore, the number of observed individuals was used as the abundance of pollinator species in each survey.

### Measuring functional floral trait

For all flowering plant species, we measured floral morphological traits that are known to be important for attracting and/or filtering pollinators: flower tube length and flower colour (Ishii et al., 2025). We measured flower corolla tube length with a calliper, and recorded flower colour based on the spectral reflectance of the flowers Spectral reflectance data were taken from Ishii et al. (2025), which examined the same study sites, for species already documented in that study, while additional measurements were made for species not previously recorded, as part of the present study (for details, see Ishii et al., 2025 and the Supplementary Materials). For all traits, we measured three (or all if there were less than three) flowers per species and used mean trait values per species. Then, we categorized flowers into three floral shape groups based on corolla tube length: open and short-tubed (< 4.5 mm), medium-tubed (4.5–9 mm), and long-tubed (> 9 mm) following previous study (Hiraiwa and Ushimaru, 2017). For flower colour, the floral reflectance spectrum of each species was translated into coordinates within the two-dimensional representations of bee and fly colour vision systems: bee colour hexagon model by Chittka (1992) and fly vision model by Troje (1993) (Fig. S1). For the bee colour system, we categorised flowers into two groups using the bee colour hexagon model: bee-non-blue (including bee-green, bee-green-blue, bee-UV-green, bee-UV) and bee-blue (including bee-blue and bee-UV-blue) flowers following Ishii et al. (2019, 2025). For the fly visual system, chromatic representation is based on the relative excitations of the two p-type and two y-type receptors (Troje 1993), and we classified flowers into two groups: fly-y-[fly-yellow (p-/y-), fly-purple (p+/y-)] and fly-y+[fly-blue (p-/y+) and fly-UV (p+/y+)] flowers (Ishii et al., 2025). Plant species in bee-non-blue and fly-y+ flowers, and those in bee-blue and fly-y+ flowers roughly correspond to each other (Fig. S1).

For each pollinator species, we measured proboscis length which influences their interactions with flowers corresponding to their tube length (Stang et al., 2007; Hiraiwa and Ushimaru, 2025). We measured four (or all if there were less than four) individuals (pollinators) per species and used mean trait values per species. Then, we categorised pollinators into three groups based on proboscis length for each site: short-tongued (< 4.5 mm), medium-tongued (4.5–9 mm), and long-tongued (> 9 mm) (Hiraiwa and Ushimaru, 2017). Moreover, pollinators were categorised into five taxonomic groups: bee, butterfly, fly, hoverfly, and others (Hirayama et al., 2025), each with distinct visual systems and colour preferences (Briscoe and Chittka, 2001; Fenster et al., 2004).

### Calculation of plant generality

To examine the relationship between floral traits and generality for each flower species (plant generality), we calculated the following three metrics. First, species generality (hereafter SG), which is defined as 1 - d’, where d’ is the species-level specialisation that measures the extent of specialisation of a flower species based on its interaction frequencies with pollinators in the network (Bluthgen et al., 2006). SG varies from 0 (completely specialist plants visited by a few pollinator species) to 1 (extremely generalist plants visited by a large number of pollinator species). Second, we calculated functional generality (FG) as the functional dispersion (FD_is_) of the proboscis lengths of interacting pollinators for each plant species, weighted by their interaction frequencies. Lower FG values indicate functional specialisation (the plants interact with pollinators having similar proboscis length), while higher FG values indicate functional generalisation (the plants interact with pollinators having different proboscis length). Third, taxonomic generality (TG), defined as FD_is_ of pollinator taxonomic groups (bees, butterflies, hoverflies, other flies and others). Low TG indicates taxonomic specialisation (visited by pollinators within few taxonomic groups), while high TG indicates taxonomic generalisation (visited by pollinators across many taxonomic groups). To avoid the effects of low sampling sizes in the plant generality index calculations, we included only plant species with at least five visits per study site, ensuring sufficient data to detect differences in their generality (Gomez-Martínez et al., 2022). SG and FD_is_ were calculated using the bipartite (Dormann et al., 2008) and FD (Laliberté & Legendre, 2010) packages, respectively, in R (R Development Core Team).

### Community-wide pollination success of plant species

We investigated pollen receipt on the stigmas of 6 and 14 species with various tube lengths and colours in 2021 and 2022, respectively (total 15 species, Table S1). These plant species were relatively widely distributed in both old and restored grasslands (Hirayama et al., 2025). During the three seasonal surveys (late June, late July, and mid-August) in 2021 and four seasonal surveys (late June, late July, mid-August, and early September) in 2022, we collected a single stigma from an open-pollinated, withered flower from each of the 20 individuals (or more than 5 individuals if there were <20) for each species at each site in each survey period. Sites where individuals of target species were less than 5 within and around the study plot were not investigated for the species. Pollen receipt (numbers of conspecific and heterospecific pollen grains deposited on each sampled stigma) of each flower was examined under a microscope. Prior to subsequent analyses, the pollen receipts on each stigma were standardised for each species across all study sites using *z*-scores, that is, (observed value – mean value) / standard deviation for the species, thus, permitting inter-species comparison. In this study, we focused solely on conspecific pollen, because our previous study revealed that heterospecific pollen receipt was significantly lower than conspecific pollen receipt and had no significant relationship with the grassland age (Hirayama et al, 2025).

### Statistical analysis

*Flower richness and abundance for each trait group -* We firstly compared temporal changes in flower richness (the number of species) and abundance between floral shape or colour groups in restored grasslands. For the analyses, we used generalised linear mixed models (GLMMs with negative binomial errors and log-link function for species richness and those with Gaussian errors and identity link function for flower abundance). In each model, flower richness or abundance of total or native flowering plants was included as the response variable and the grassland age, floral shape or colour group (open and short-tubed/medium-tubed/long-tubed or bee-non-blue/bee-blue or fly-y-/fly-y+), the interaction between them and the spatial auto-covariates as the explanatory variables alongside with independent random terms of study year identity and the number of surveys per site (four/five per year). We constructed these models separately for the shape group (long/middle/short) and the colour groups (bee-non-blue/bee-blue or fly-y-/fly-y+). We calculated the spatial auto-covariates from the latitude and longitude measurements of all sites to consider the effects of spatial autocorrelation on statistical analyses (Dormann et al., 2007) because the study sites were not randomly distributed within the study area. Study year identity and survey time were included as independent random terms to avoid the effects of temporal pseudo-replication and difference in sampling effort. We did not include site identity as a random term because the spatial auto-covariates already control unexamined site specific effects.

We also compared flower richness and abundance of each floral shape or colour group between grassland types (restored/old), we constructed a GLMM with each flower richness or abundance parameter as the response variable and six or four flower-grassland categories (3 shape groups × 2 grassland types, 2 bee-colour groups × 2 grassland types or 2 fly-colour groups × 2 grassland types) were included as explanatory variables, respectively. Study year identity and survey time and spatial auto-covariate were included as random terms and the explanatory variable in the models. Subsequently, we conducted a multiple comparison among flower-grassland categories by Tukey’s method using the ‘multcomp’ package (Hothorn et al., 2008).

*Relationships between floral trait composition and pollinator composition*- To examine the relationship between floral traits and pollinator composition, we first conducted principal component analysis (PCA) using data of the relative flower abundance of each floral trait group. Flower grouping was based on combinations of floral shape and bee colour group, and floral shape and fly colour group. Hereafter we refer these as “flower PCs” with bee and fly colour groupings. Second, we also conducted PCA for pollinator composition using data of the relative abundances for the trait and taxonomic groups of pollinators: for example, short-tongued flies, medium-tongued hoverflies, long-tongued bees. Hereafter we refer to these as “pollinator PCs”. Finally, we compared pollinator PCs and flower PCs with bee and fly colour groupings, respectively, using Pearson’s correlation coefficient with Bonferroni correction for multiple comparisons. We also conducted the same analyses for native plant species and their pollinators.

*Plant generality* - To examine how generality of each floral shape or colour group have changed during the grassland restoration duration and differences in temporal change pattern among the group in restored grasslands, we constructed a GLMM in which the response variable was each plant generality metric (SG or FG or TG) of each flower species and the explanatory variables were grassland age, floral group (shape, bee-colour or

fly-colour group), the interaction between them and the spatial auto-covariates. We included study year identity and the survey time as independent random terms in each model. We conducted GLMMs for total and native plant species seperately.

*Pollination success-* We compared temporal changes in community-wide pollen receipt during grassland restoration among floral trait groups using GLMMs (Gaussian errors and identity link function). In the GLMM, the standardised conspecific pollen number on each stigma for each species for each site were included as the response whereas grassland age, floral trait (shape or colour) group and their interaction were included as the explanatory variables. For the 15 study species, the bee colour grouping (bee-non-blue/bee-blue) was identical to the fly colour grouping (fly-y-/fly-y+), respectively. In each GLMM, we incorporated spatial auto-covariates as a covariate, and study year and survey month identities as nested random terms in the models.

We also examined the differences in the relationships between pollen receipt and each plant generality metric among floral trait groups with GLMMs. We incorporated the standardised conspecific pollen number on each stigma for each species for each site as the response variables and generality metrics of each species, floral trait group and their interaction as the explanatory variables. In the above generality-grassland age relationship analysis, we found no significant differences in temporal changes in the TD among floral trait groups. Therefore, in this analysis, we only examined SG and FG as generality metrics. We incorporated visiting frequency (visits/ flower abundance (m²)) for each species, spatial auto-covariates as a covariate, and study year and survey month identities as nested random terms in the models.

## RESULTS

In the main text, we primarily focus on the results of native species, while corresponding analytical results for total species are provided in the supplementary materials (Table S3). This emphasis aligns with our primary research objective of understanding the recovery process of native plant communities in restored grasslands. Notably, the results were largely consistent for both total and native flowers (Table S3).

### Species richness and flower abundance of each floral traits

Native flower richness significantly differed among the floral shape groups, such that open/short-tubed flowers exhibited higher richness than middle- and long-tubed flowers in restored grasslands (Fig. 1a, Table S2). Similarly, the richness of bee-non-blue and fly-y-minus flowers was higher than that of bee-blue and fly-y-plus flowers in restored grasslands (Fig. 1b, c, Table S2). The temporal change patterns in flower richness significantly differed between floral shape groups, such that open/short-tubed flowers exhibited much shallower regression slopes than long-tubed flowers (Fig. 1a, Table S2). Meanwhile, there were no significant differences in temporal change patterns in bee- and fly-colour group, respectively (Fig. 1c, Table S2). We also found that the mean richness of long-tubed and fly-y+ flowers remained lower in restored grasslands than in old grasslands (Fig. S2). Difference in these patterns in flower abundance among trait groups showed similar but more conspicuous trends (Fig. 1d, e, f, Fig. S2, Table S2). Specifically, bee- and fly-colour groups showed significantly different temporal change patterns in each group (Fig. 1e, f, Table S2), unlike species richness. These results indicate that long-tubed, bee-blue, and fly-y+ flowers take more time to recover compared to open/short-tubed, bee-non-blue and fly-y- flowers in restored grasslands, respectively. These trends were almost consistent for both total and native flowers (Table S3).

**Figure 1.**
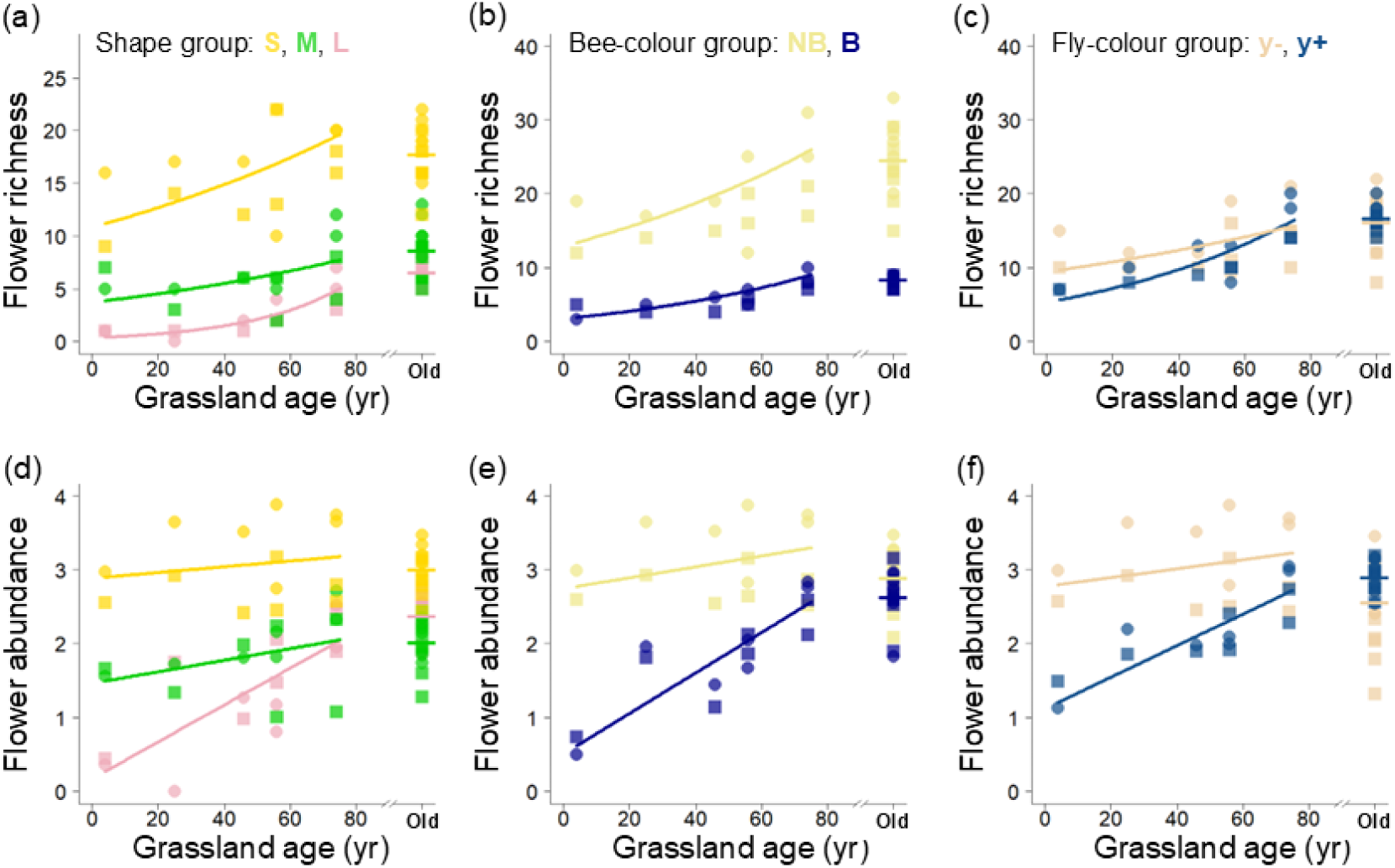
Temporal changes in native flower richness and abundance of each shape group (a and d, respectively: yellow, short-tubed, S; green, medium-tubed, M; pink, long-tubed, L), bee-colour group (b and e, respectively: bee-non-blue, NB; bee-blue, B) and fly-colour group (c and f, respectively: fly-y-, y-; fly-y+, y+) along grassland age (years since restoration started, yr) in the restored grasslands, and these parameters in old grasslands with horizontal lines indicating their means as references: data of 2021 and 2022 are shown as squares and circles. The solid lines represent regressions estimated from the GLMMs (Table S2).

### Trait composition in flower and pollinator

For PCA using the floral shape-bee colour group dataset [short-tubed bee-non-blue (S-NB), short-tubed bee-blue (S-B), middle-tubed bee-non-blue (M-NB), middle-tubed bee-blue (M-B), long-tubed bee-non-blue (L-NB), long-tubed bee-blue (L-B)], PC1 (45.96%) and PC2 (19.37%) explained 65.33% of the total variance. PC1 was positively correlated with the relative abundances of L-NB, M-NB and all bee-blue (S-B, M-B, and L-B) flowers and negatively correlated with that of S-NB flowers (Fig. 2a). PC2 was positively correlated with the relative abundances of S-B flowers, and negatively correlated with those of M-NB and L-B flowers (Fig. 2a). For PCA using the shape-fly colour group dataset [short-tubed fly-y- (S-y-), short-tubed fly-y+ (S-y+), middle-tubed fly-y- (M-y-), middle-tubed fly-y+ (M-y+), long-tubed fly-y- (L-y-), long-tubed fly-y+ (L-y+)], PC1 (44.07%) was positively correlated with the relative abundances of M-y- and all fly-y+ (S-y+, M-y+, and L-y+) flowers and negatively correlated with that of S-y- flowers (Fig. 2b). PC2 (25.38%) was positively correlated with the relative abundance of L (L-y- and L-y+) and S-y+ flowers, and negatively with those of medium-tubed (M-y- and M-y+) and S-y- flowers (Fig. 2b).

**Figure 2.**
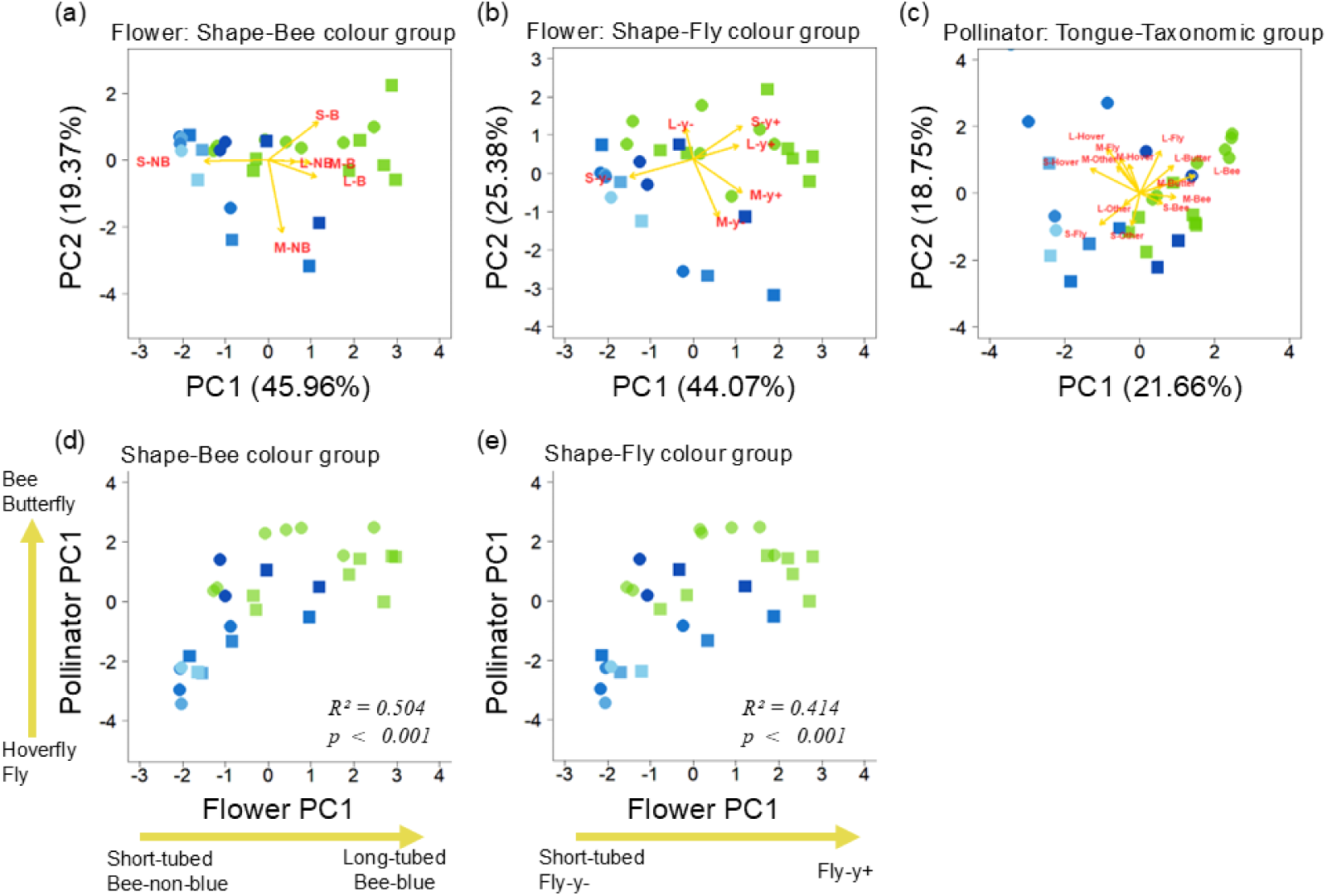
PCA biplots of flower trait groups (a: 3 shape groups × 2 bee colour groups), (b: 3 shape groups × 2 fly colour groups) and pollinator trait and taxonomic groups (c: 3 proboscis groups × 5 taxonomic groups) in restored (blue, the younger had a lighter colour) and reference old grasslands (green). Relationships between flower PC1 and pollinator PC1 and PC2 (d, flower PCA by shape-bee colour group; e, by shape-fly colour group) in the restored grasslands and old grasslands. Data of 2021 and 2022 are shown as squares and circles.

For pollinators, we found that the first two PCA axes explained 40.38 % of the total variance (PC1: 21.66%, PC2: 18.75%). PC1 was positively correlated with the relative abundances of bees, butterflies and long-tongued flies, and negatively correlated with those of hoverflies, flies and others (Fig. 2c). PC2 was positively correlated with the relative abundance of longer-tongued species for each taxonomic group, and negatively correlated with those of shorter-tongued bees, flies, and other insects (Fig. 2c).

Pollinator PC1 exhibited significant positive relationships with flower PC1s with both bee and fly colour groupings (Fig. 2d, e; bee colour, *r* = 0.504, *p <* 0.001; fly colour, *r* = 0.414, *p <* 0.001), indicating that the proportion of bees and butterflies increased with the proportion of long-tubed and bee-blue/fly-y+ flowers within a network. Meanwhile, pollinator PC2 had no significant relationships with flower PCs in both bee and fly colour grouping (Table S4). The same trends were observed for total flowers (Fig. S3, Table S4).

### Generality of each floral trait

In young restored grasslands, species generality (SG) of short-tubed flowers was significantly higher than that of long-tubed flowers (Fig. 3a, Table S2). The slopes of grassland age-SG regressions significantly differed between short-/medium-tubed and long-tubed flowers (Fig. 3a, Table S2). For colour groups, bee-non-blue and fly-y- flowers had higher SGs than bee-blue and fly-y+ flowers, respectively (Fig. 3b, c, Table S2).

**Figure 3.**
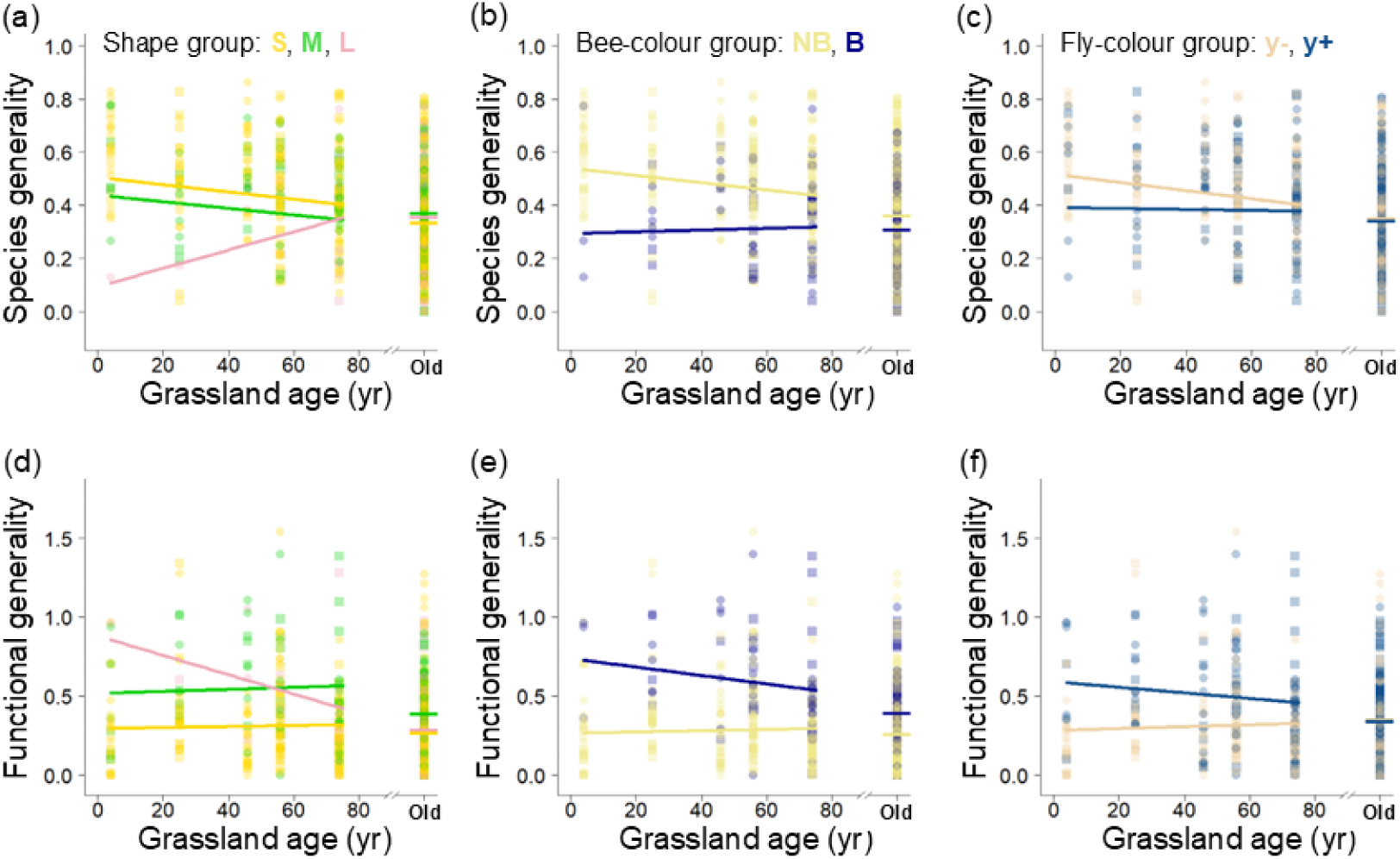
Temporal changes in species generality and functional generality of native plant species of each shape group (a and d, respectively: yellow, short-tubed, S; green, middle-tubed, M; pink, long-tubed, L), bee-colour group (b and e, respectively: bee-non-blue, NB; bee-blue, B) and fly-colour group (c and f, respectively: fly-y-, y-; fly-y+, y+) along grassland age (years since restoration started, yr) in the restored grasslands, and these parameters in old grasslands with horizontal lines indicating their means as references: data of 2021 and 2022 are shown as squares and circles. The solid lines represent regressions estimated from the GLMMs (Table S2).

Functional generalities (FGs) in middle-/long-tubed, bee-blue and fly-y+ flowers were higher than those in short-tubed, bee-non-blue and fly-y- flowers, respectively, in young restored grasslands (Fig. 3d, e, f, Table S2). Moreover, the interactions between grassland age and floral shape groups had significant effects, such that FG of long-tubed flowers decreased with grassland age, whereas that of short-/medium-tubed flowers did not change across grassland age in restored grasslands (Fig. 3d, Table S2). Meanwhile, the interactions between grassland age and floral colour groups showed no significant effects on FGs (Fig. 3e, f, Table S2). Taxonomic generalities (TGs) showed no significant differences among all floral trait groups and exhibited no temporal changes across any of these groups (Table S2). These trends were almost consistent for both total and native flowers (Table S3).

### Pollination success of each floral trait

Compared to short-/medium-tubed flowers, long-tubed flowers received less conspecific pollen in young restored grasslands, and only that of middle-tubed flowers increased with grassland age significantly greater than short-/long-tubed flowers (Fig. 4a, Table S2). Meanwhile, pollen receipt of bee-non-blue (fly-y-) flowers were significantly more than that of bee-blue (fly-y+) flowers in young restored grasslands, and both increased similarly with grassland age (Fig. 4d, Table S2). Conspecific pollen receipts of short-tubed and bee-non-blue (fly-y-) flowers did not vary with plant SG and FG, whereas those of medium-/long-tubed and bee-blue (fly-y+) flowers decreased with plant SG and/or FG (Fig. 4b, c, e, f, Table S2).

**Figure 4.**
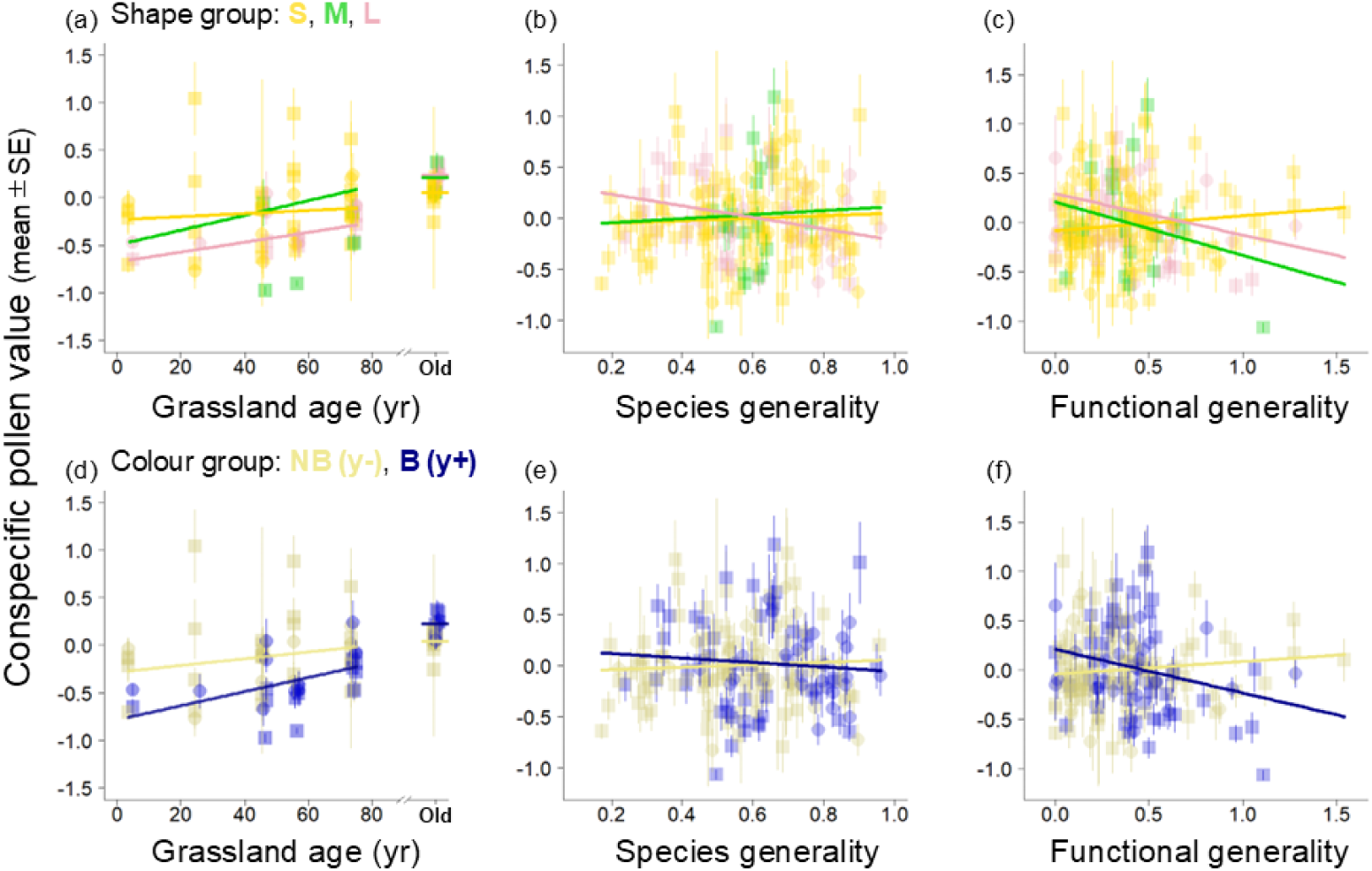
Temporal changes in conspecific pollen value of each shape group (a: yellow, short-tubed, S; green, middle-tubed, M; pink, long-tubed, L) and colour group (d: bee-non-blue, NB; bee-blue, B) in the restored grasslands, and these parameters in old grasslands with horizontal lines indicating their means as references: data of 2021 and 2022 are shown as squares and circles. Relationships between conspecific pollen value and both species and functional generality (b, c, e and f). Bee colour groups (NB/B) were identical to fly colour groups (y-/y+), respectively (Table S1). The solid lines represent regressions estimated from the GLMMs (Table S2).

## DISCUSSION

Trait-based assembly perspective in relation to mutualistic trophic interactions would advance a prediction of community reassembly pattern in ecological restoration. By focusing on plant-pollinator interactions, we provide empirical evidence that floral tube length and colour were important traits determining the plant reassembly process through their interactions with pollinators during the long-term grassland restoration. We further found contrasting temporal change patterns of species and functional generality for plant species with different traits in restored grasslands, and their effects on their pollination success in this plant community reassembly process. Overall, our findings support our prediction that species with generalist-adapting floral traits (open/short tubes and bee-non-blue/fly-y-colours) have recovered more quickly than those with specialist flowers with long-tubes and/or bee-blue/fly-y+ colours in restored grasslands. Moreover, higher plant functional specialisation (lower functional generality) rather than species specialisation promoted higher pollination success of such species with specialist flowers in older restored grasslands, suggesting their importance of functional specialisation in the recovery of the plants. Below we discuss our findings in more detail in the light of trait-based reassembly of plant community in long-term grassland restoration.

We found that the richness and abundance of all floral trait groups increased with the age of restored grassland, although the recovery patterns significantly differed among the trait groups. As expected, plants with open/short-tubed and bee-non-blue/fly-y- (white and yellow to human eyes) flowers were more likely to reestablish rapidly in young restored grasslands (Fig. 1). In contrast, plants with long-tubed and bee-blue/fly-y+ (blue, purple and red to human eyes) flowers took longer to recover, and their richness and abundance had not reached the levels observed in old grasslands even after 75 years. Consequently, younger restored grasslands were more dominated by flowers with open/short tubes and white or yellow colours. Pollinator composition is typically linked to floral shape and colour compositions (Stang et al., 2007; Ishii et al., 2019, 2025). We found temporal shifts in pollinator community composition from those dominated by shorter-tongued dipterans to those with more abundance of longer-tongued bees, mainly bumblebees (Hirayama et al., 2025; this study). These pollinator shifts corresponded to the shifts in floral trait composition in restored grasslands, being consistent with previous studies (Stang et al., 2007; Ishii et al., 2019; Ishii et al., 2025). Such long-term dynamics of plant and pollinator communities during the long-term restoration periods may influence visitation patterns to individual plant species and the overall structure of plant-pollinator networks.

In this study, we hypothesised that floral trait reassembly patterns are well explained by considering the degree of generality for each trait group. Supporting our predictions, flowers with traits that attract a broader range of pollinators (generalist flowers) have recovered more quickly in restored grasslands. Open/short-tubed and white/yellow flowers had higher species generality than long-tubed and blue/purple flowers in young restored grasslands. Generalist plant species are more resilient to more limited pollinator fauna due to their high potential to interact with a wide variety of partners (Astegiano et al., 2015; Proesmans et al., 2016). On the other hand, the low functional generality for these generalist flowers was observed in young restored grasslands, being likely due to that they were visited by diverse short-tongued pollinators. Furthermore, the conspecific pollen receipts of open/short-tubed and white/yellow flowers were higher throughout the grassland restoration period, regardless of species or functional generality. Thus, generalist plant species have consistently maintained high pollination success, outweighing the risk of interspecific pollination by generalist pollinators. This further strengthens the hypothesised causal pathway linking floral traits, pollinator generality, and plant community reassembly in the grassland restoration duration.

In contrast, long-tubed and blue/purple flowers exhibited lower species generality and higher functional generality in younger restored grasslands. High functional generality observed in young restored grasslands likely contributed to this slow recovery of these flowers through low pollination success caused by trait mismatching between these flowers and available pollinators (Hiraiwa and Ushimaru, 2024). Moreover, during the restoration process, species generality for these specialist flowers increased while their functional generality decreased, suggesting that such flowers have become visited by various long-tongued pollinators, likely several long-tongued bee species (Fig. S4, 5, 6). Thus, the observed temporal change patterns in these metrics for these specialist flowers indicate that functional generality was particularly critical for their reproductive success, although both lower species and functional generality enhanced conspecific pollen receipt (Fig. 4). The reduction in functional generality (increase in functional specialisation) may compensate for the potential negative effects of higher species generality by ensuring more effective pollen transfer through promoting trait-matching with specialized pollinators. Thus, functional specialisation rather than species specialisation appears pivotal for the reproductive success of plant species with specialist flowers, consistent with the previous findings (Hiraiwa and Ushimaru, 2017, 2024).

Interestingly, the levels of both species and functional generality of plants with different floral traits were more similar in reference old grasslands than those in young and old restored grasslands (Fig. 3). This may suggest that in mature grasslands with high species and functional diversity of plants and pollinators, resource competitions among pollinators with diverse functional traits facilitated functional niche partitioning among them (Hiraiwa and Ushimaru, 2017, 2024).

The differences in reassembly patterns between generalist- and specialist-pollinated plant species should influence temporal transition of pollination network structure during grassland restoration. In our previous study conducted in the same sites, we found that the pollination networks were highly generalised in young restored grasslands, but as plant species recovery progressed over decades, the networks gradually increased the degree of specialisation (Hirayama et al., 2025). This observed transitional pattern is likely attributed to the quick recovery of generalist flowers at the early stages of restoration, followed by the delayed and slow recovery of plants with specialist flowers. Increases in functional diversity of flowers also likely promoted functional niche partitioning among pollinators in old restored grasslands, resulting in functional generality for each floral trait group approaching that in old grasslands. This, in turn, promoted higher network level specialisation in old restored grasslands compared to young restored grasslands.

In contrast to our findings, a pollination network assembly in alpine grasslands established on glacier-retreated areas is reported to progress towards generalisation (Albrecht et al., 2010). This may be explained by the floral trait associations with the area-specific pollinator communities. Alpine communities are typically characterized by small dipteran pollinator dominance (Albrecht et al., 2010; Kudo et al., 2024; Ishii et al., 2025), where such generalist pollinators enhance the recovery of diverse plant species with generalist floral traits. Thus, trait-based plant (re)assembly patterns and consequent network structures in mature vegetation would differ depending on potential pollinator compositions of the target area (Ishii et al., 2025). Overall, focusing on floral traits and their interactions with pollinators, we can predict temporal transitional patterns in flower and pollinator composition, network structure, and pollination function during community reassembly in restored grasslands.

## Conclusion

Our findings provide novel insights into the role of floral traits in shaping temporal patterns in plant and pollinator reassembly and pollination network dynamics during grassland restoration. Generalist floral traits such as open/short corolla tubes and bee-non-blue/fly-y-colours exhibited faster recovery at the early restoration stages, being maintained by interactions with a wide range of short-tongued pollinators. As restoration progresses, the recovery of functional matching between specialist flower species and long-tongued bees contributed to increased network specialisation, suggesting the co-reassembly of plant and pollinator communities. While the causal relationship between specialist plants and specialist pollinators remains unclear, recovery of their reciprocal interactions likely contributes significantly to overall community reassembly. However, examining a limited number of floral and pollinator traits (tube length and colour, and tongue length) may not fully capture the whole picture of complex plant-pollinator interactions within restored communities. Future research exploring additional floral traits (such as flowering phenology, floral scent, and pollen and nectar production) and other types of terrestrial vegetation are warranted. This would build a more comprehensive understanding of the plant and pollinator reassembly process, which can improve the effectiveness and efficiency of ecosystem restorations.

## Acknowledgements

We thank the land owners of our study sites, which are ‘Sugadaira Bokujo’, ‘Hatsune-kan’, ‘Imai-kan’, and ‘Jozan-kan’; and the management companies, ‘Sugadaira Pine beak ski’, ‘Sugadaira Ski House Co., Ltd.’, ‘Oku Davos Snow Park’, and ‘HARE Sugadaira-Kogen Snow Resort’ for permitting to conduct field surveys. We thank Yaida A. Yuki for helping to investigate grassland history in studied grasslands. We greatly thank for Kouki Okusawa getting floral trait data in this study. This work was supported by the Sasakawa Scientific Research Grant from The Japan Science Society, the fund of Nagano Prefecture to promote scientific activity to G.S.H., and a Grant in Aid for Scientific Research Programs (KAKENHI no. 19H03303 and 22K06400) from the Japan Society for the Promotion of Science to A.U. and H.S.I This research was also performed by the Environment Research and Technology Development Fund (JPMEERF20234005) of the Environmental Restoration and Conservation Agency provided by Ministry of the Environment of Japan.

## Author contribution

Gaku S. Hirayama, Tanaka Kenta, and Atushi Ushimaru conceived the study concept and Gaku S. Hirayama and Atushi Ushimaru developed the methodology studying community-wide pollination networks; Gaku S. Hirayama basically collected and analysed the whole data; Hiroshi S. Ishii, Taisei W. Ueno and Atushi Ushimaru collected flower trait data; Tanaka Kenta also helped Gaku S. Hirayama to conduct the pollination experiment; Gaku S Hirayama wrote first draft together with Atushi Ushimaru. All authors contributed critically to the drafts and gave final approval for publication.

