## Supplement files for "Generalist flowers recover rapidly in grassland restoration: The importance of floral traits in community reassembly"

**Supplement materials**

**Flower tube and spectral reflectance of flowers**

We measured flower tube length using a digital calliper (CD-15PS; Mitutoyo, Kawasaki, Japan). Flower tubes were defined as tubular structures formed by the receptacle, calyx, and corolla or a combination of these organs. In the absence of a tube, the tube length was regarded as 0 mm.

We measured the spectral reflectance using a charge-coupled device spectrometer (BRC112; B&W Tek, Newark, DE), Y-shaped fibre optic cable (FRP-400-0.22-1.5-UV; B&W Tek), a white standard (SRR; B&W Tek, or freshly pressed pellet of dry BaSO₄), and deuterium-tungsten light sources (BDS130; B&W Tek). Measurements were taken at 1 nm wavelength intervals from 300 to 700 nm. For flowers with multiple colours, we targeted the most conspicuous region: the banner petal for a *Fabaceae* species, the ray florets in composite capitula containing both disc and ray florets, and the outer region in actinomorphic flowers.

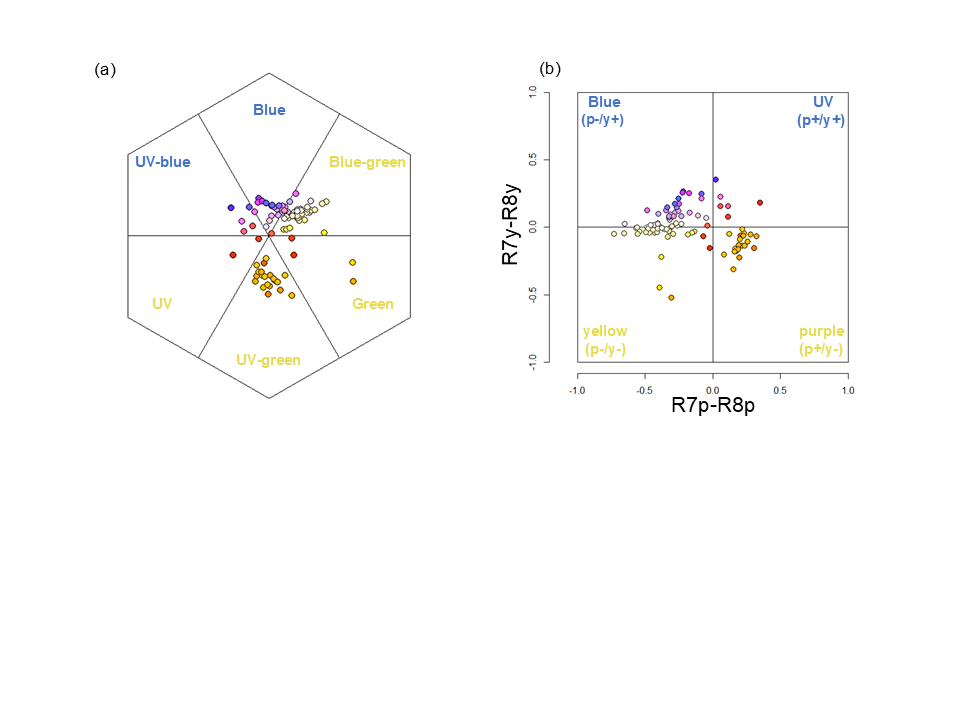

**Figure S1.** Locations of floral colours of the study species within the bee color hexagon (a, Chittka, 1992) and the fly visual model (b, Troje, 1993). Colours of the symbols represent the chromaticity of the flowers as perceived by the human eye.

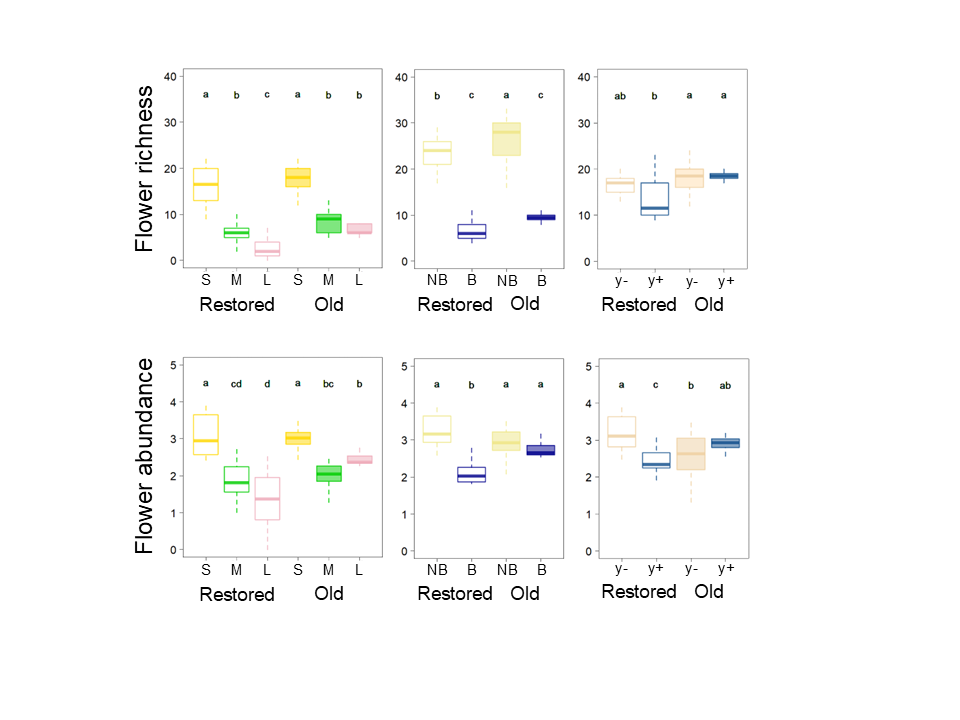
**Figure S2.** Comparison of richness and abundance of native plant among shape (yellow, S; green, M; pink, L) or colour (cream, bee-non-blue, NB; blue, bee-blue, B; light brown, fly-y-, y-; light blue, fly-y+, y+) groups between old and restored grasslands. The central bars indicate the medians in the boxplots, and different alphabets show significant differences (*p < 0.05*).

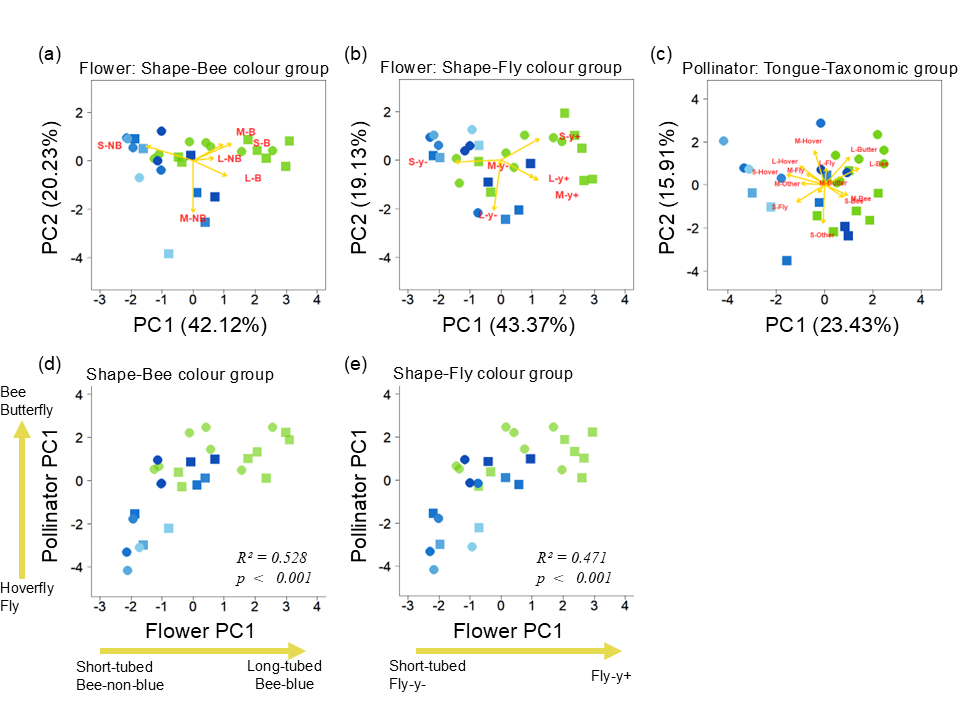
**Figure S3.** PCA biplots of flower trait groups (a: 3 shape groups × 2 bee colour groups), (b: 3 shape groups × 2 fly colour groups) and pollinator trait and taxonomic groups (c: 3 proboscis groups × 5 taxonomic groups) in restored and reference old grasslands. In shape-bee colour groups, flower PC1 (42.12%) and PC2 (20.23%) explain 62.35% of the total variance. Flower PC1 was positively correlated with the relative abundances of long-tubed bee-non-blue (L-NB) and all bee-blue (B) flowers and negatively correlated with that of short-tubed bee-non-blue (S-NB) flowers. Flower PC2 was positively correlated with the relative abundance of short-/medium-tubed bee-non-blue (S-NB and M-NB) flowers and short-tubed bee-blue (S-B) flowers, and negatively with long-tubed bee-blue (L-B) and M-NB flowers. Similarly, in shape-fly colour groups, flower PC1 (43.37%) was positively correlated with the relative abundances of all fly-y+ flowers and negatively correlated with that of short-tubed fly-y- flowers. Flower PC2 was positively correlated with the relative abundance of short-tubed fly-y+ (S-y+) flowers, and negatively with medium-/long-tubed flowers (M-y-, M-y+, L-y-, L-y+). Pollinator PC1 (23.43%) and PC2 (15.91%) explain 39.34 % of the total variance. Pollinator PC1 was positively correlated with relative abundances of bees and butterflies, and negatively with those of hoverflies and other flies. Pollinator PC2 was positively correlated with the relative abundance of longer tongued species of all taxonomic groups, and negatively with shorter-tongued bees, flies and others. Relationships between flower PC1 and pollinator PC1 and PC2 (d, flower PCA by shape-bee colour group; e, by shape-fly colour group) in the restored grasslands (blue, the younger had a lighter colour) and old grasslands (green). Data of 2021 and 2022 are shown as squares and circles. The solid lines represent significant regressions from the GLMMs (Table S4).

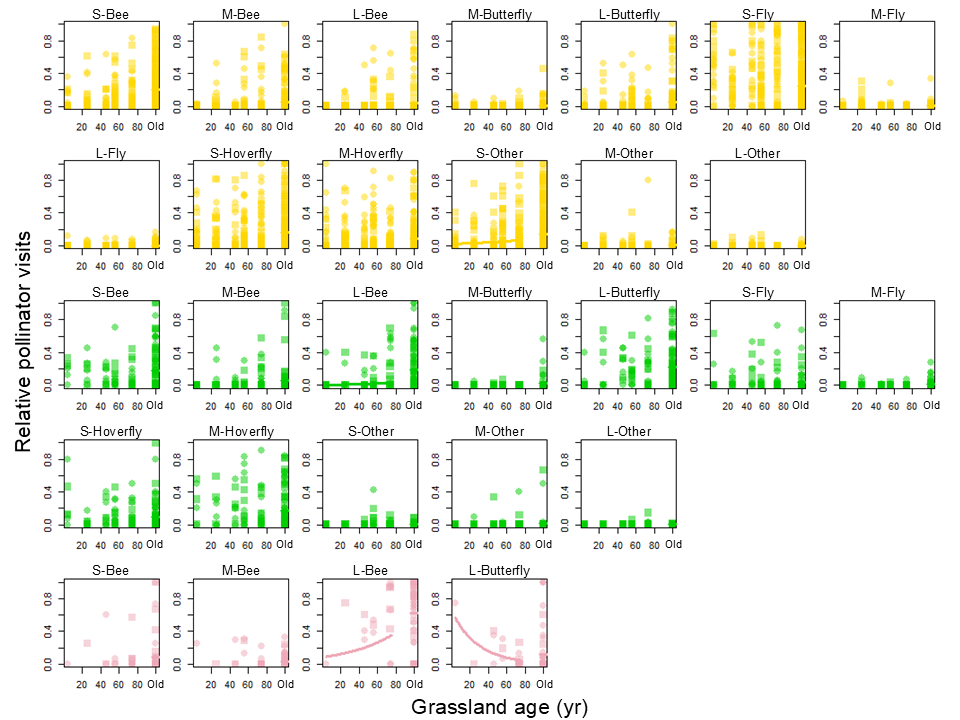

**Figure S4.** Temporal changes in the relative visits by each pollinator group (3 proboscis groups × 5 taxonomic groups) to total visits in different floral shape groups (yellow, S; green, M; pink, L) of native plant species along grassland age in restored grasslands, as well as these pollinator parameters in old grasslands: data of 2021 and 2022 are shown as squares and circles, respectively. Pollinator groups with no or rare visits (< 5 visits) to the given floral shape groups were not included in the statistical analyses and this figure. The solid lines indicate significant regressions estimated from GLMMs (*p < 0.05*).

.

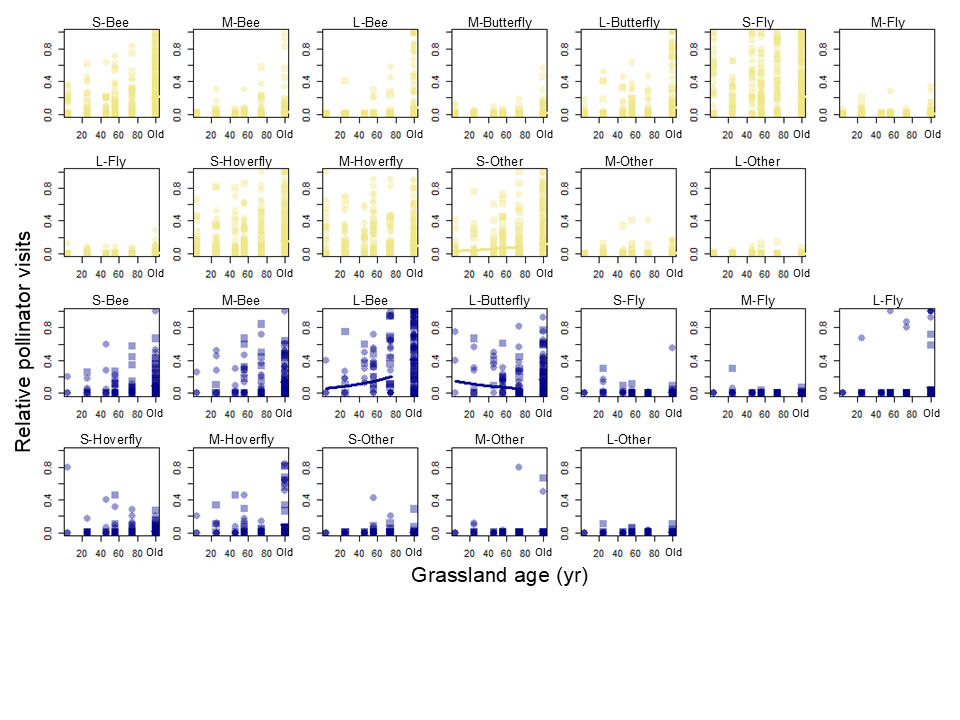

**Figure S5.** Temporal changes in the relative visits by each pollinator group (3 proboscis groups × 5 taxonomic groups) to total visits in different floral bee-color groups (cream, NB; blue, B) of native plant species along grassland age in restored grasslands, as well as these pollinator parameters in old grasslands: data of 2021 and 2022 are shown as squares and circles, respectively. Pollinator groups with no or rare visits (< 5 visits) to the given floral shape groups were not included in the statistical analyses and this figure. The solid lines indicate significant regressions estimated from GLMMs (*p < 0.05*).

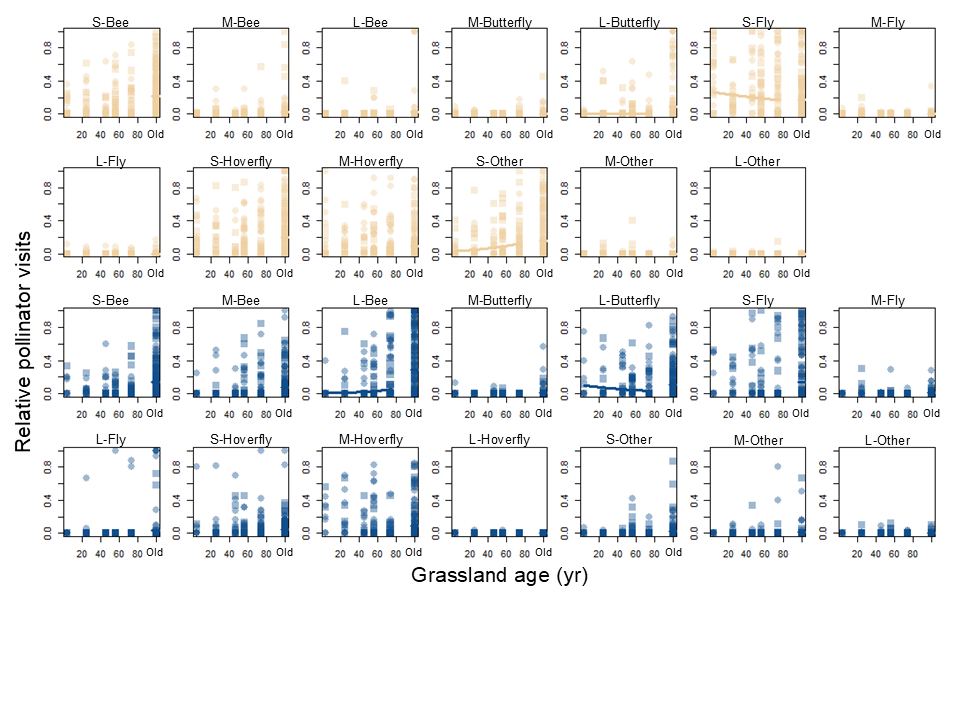

**Figure S6.** Temporal changes in the relative visits by each pollinator group (3 proboscis groups × 5 taxonomic groups) to total visits in different floral fly-color groups (light brown, y-; light blue, y+) of native plant species along grassland age in restored grasslands, as well as these pollinator parameters in old grasslands: data of 2021 and 2022 are shown as squares and circles, respectively. Pollinator groups with no or rare visits (< 5 visits) to the given floral shape groups were not included in the statistical analyses and this figure. The solid lines indicate significant regressions estimated from GLMMs (*p < 0.05*).

**Table S1.** Information of plant species used in pollination (pollen) surveys: flowering phenology, floral shape group, and colour groups were shown.

| Family | Species | Flowering phenology | Shape group | Colour group (bee-, fly-) |
| --- | --- | --- | --- | --- |
| Asteraceae | *Ixeridium dentatum* | Jun.-Jul. | S | NB, y- |
| Asteraceae | *Eupatorium glehni* | Jul.-Aug. | S | NB, y- |
| Asteraceae | *Picris hieracioides* | Aug.-Sep. | M | NB, y- |
| Asteraceae | *Cirsium oligophyllum* | Aug.-Sep. | M | B, y+ |
| Campanulaceae | *Platycodon glandiflorus* | Aug.-Sep. | S | B, y+ |
| Fabaceae | *Lespedeza bicolor* | Aug.-Sep. | M | B, y+ |
| Iridaceae | *Iris sanguinea* | Jun.-Jul. | L | B, y+ |
| Lamiaceae | *Prunella vulgaris* | July | L | B, y+ |
| Lamiaceae | *Thymus quinquecostatus* | July | M | NB, y- |
| Liliaceae | *Polygonatum odoratum* | May-Jun. | L | NB, y- |
| Primulaceae | *Lysimachia clethroides* | July | S | NB, y- |
| Ranunculaceae | *Ranunculus japonicus* | May-Jun. | S | NB, y- |
| Rosaceae | *Agrimonia pilosa* | Jul.-Aug. | S | NB, y- |
| Rosaceae | *Sanguisorba officinalis* | Aug.-Sep. | S | NB, y- |
| Rubiaceae | *Galium verum* | July | S | NB, y- |

**Table S2** Results of generalized linear mixed model analyses for richness, abundance, plant generality and conspecific pollen receipt for native plant species. Short-tubed, bee-non-blue and fly-y-minus (fly-y-) flowers were used as the baselines for floral shape and bee- and fly- colour groups, respectively. Significant effects are indicated in bold (*P <* 0.05).

| **Response variables** | | **Explanatory variable** | **Coefficients** | | **z** | **P** |
| --- | --- | --- | --- | --- | --- | --- |
|  |  |  | **Estimate** | **SE** |  |  |
| **Shape** | Flower richness | **Grassland age (Age)** | 0.008 | 0.003 | 2.376 | **0.018** |
|  |  | **Medium-tubed (Mt)** | -1.066 | 0.320 | -3.334 | **< 0.001** |
|  |  | **Long-tubed (Lt)** | -3.376 | 0.662 | -5.104 | **< 0.001** |
|  |  | Age * Mt | 0.002 | 0.006 | 0.316 | 0.752 |
|  |  | **Age * Lt** | 0.027 | 0.011 | 2.549 | **0.011** |
|  |  | Spatial autocovariate | 0.000 | 0.000 | -1.808 | 0.071 |
|  |  | **Intercept** | 2.498 | 0.173 | 14.417 | **< 0.001** |
|  | Flower abundance | Age | 0.004 | 0.006 | 0.68 | 0.496 |
|  |  | **Mt** | -1.424 | 0.397 | -3.588 | **< 0.001** |
|  |  | **Lt** | -2.714 | 0.397 | -6.839 | **< 0.001** |
|  |  | Age * Mt | 0.004 | 0.007 | 0.532 | 0.595 |
|  |  | **Age * Lt** | 0.021 | 0.007 | 2.834 | **0.005** |
|  |  | Spatial autocovariate | 0.000 | 0.000 | -0.048 | 0.962 |
|  |  | **Intercept** | 2.892 | 0.284 | 10.2 | **< 0.001** |
| **Colour (bee)** | Flower richness | **Age** | 0.009 | 0.003 | 2.936 | **0.003** |
|  |  | **Bee-blue (Bb)** | -1.423 | 0.338 | -4.206 | **< 0.001** |
|  |  | Age * Bb | 0.005 | 0.006 | 0.830 | 0.406 |
|  |  | Spatial autocovariate | 0.000 | 0.000 | -1.775 | 0.076 |
|  |  | **Intercept** | 2.580 | 0.165 | 15.631 | **< 0.001** |
|  | Flower abundance | Age | 0.007 | 0.004 | 1.650 | 0.099 |
|  |  | **Bb** | -2.237 | 0.300 | -7.447 | **< 0.001** |
|  |  | **Age * Bb** | 0.020 | 0.006 | 3.559 | **< 0.001** |
|  |  | Spatial autocovariate | 0.000 | 0.000 | -1.827 | 0.068 |
|  |  | **Intercept** | 2.902 | 0.250 | 11.626 | **< 0.001** |
| **Colour (fly)** | Flower richness | Age | 0.007 | 0.004 | 1.894 | 0.058 |
|  |  | **Fly-y-plus (Fy+)** | -0.572 | 0.281 | -2.037 | **0.042** |
|  |  | Age * Fy+ | 0.008 | 0.005 | 1.679 | 0.093 |
|  |  | Spatial autocovariate | 0.000 | 0.000 | -1.819 | 0.069 |
|  |  | Intercept | 2.360 | 0.187 | 12.625 | **< 0.001** |
|  | Flower abundance | Age | 0.006 | 0.004 | 1.466 | 0.143 |
|  |  | **Fy+** | -1.654 | 0.278 | -5.943 | **< 0.001** |
|  |  | **Age * Fy+** | 0.015 | 0.005 | 2.934 | **0.003** |
|  |  | Spatial autocovariate | 0.000 | 0.000 | -1.713 | 0.087 |
|  |  | **Intercept** | 2.905 | 0.244 | 11.913 | **< 0.001** |
| **Shape** | Species generality | **Age** | -0.001 | 0.001 | -2.634 | **0.008** |
|  |  | Mt | -0.067 | 0.064 | -1.047 | 0.295 |
|  |  | **Lt** | -0.411 | 0.131 | -3.145 | **0.002** |
|  |  | Age * Mt | 0.000 | 0.001 | 0.135 | 0.893 |
|  |  | **Age * Lt** | 0.005 | 0.002 | 2.831 | **0.005** |
|  |  | Spatial autocovariate | 0.000 | 0.000 | -0.379 | 0.705 |
|  |  | **Intercept** | 0.509 | 0.043 | 11.925 | **< 0.001** |
|  | Functional generality | Age | 0.000 | 0.001 | 0.356 | 0.722 |
|  |  | **Mt** | 0.222 | 0.105 | 2.11 | **0.035** |
|  |  | **Lt** | 0.585 | 0.208 | 2.806 | **0.005** |
|  |  | Age * Mt | 0.000 | 0.002 | 0.205 | 0.838 |
|  |  | **Age * Lt** | -0.006 | 0.003 | -2.209 | **0.027** |
|  |  | **Spatial autocovariate** | 0.000 | 0.000 | -2.61 | **0.009** |
|  |  | **Intercept** | 0.359 | 0.053 | 6.786 | **< 0.001** |
|  | Taxonomic generality | Age | 0.000 | 0.000 | -1.333 | 0.182 |
|  |  | Mt | -0.010 | 0.040 | -0.241 | 0.809 |
|  |  | Lt | -0.017 | 0.083 | -0.205 | 0.837 |
|  |  | Age * Mt | 0.000 | 0.001 | 0.774 | 0.439 |
|  |  | Age * Lt | -0.001 | 0.001 | -1.049 | 0.294 |
|  |  | Spatial autocovariate | 0.000 | 0.000 | -0.624 | 0.533 |
|  |  | **Intercept** | 0.261 | 0.023 | 11.291 | **< 0.001** |
| **Colour (bee)** | Species generality | **Age** | -0.001 | 0.000 | -2.762 | **0.006** |
|  |  | **Bb** | -0.246 | 0.068 | -3.608 | **< 0.001** |
|  |  | Age * Bb | 0.002 | 0.001 | 1.668 | 0.095 |
|  |  | Spatial autocovariate | 0.000 | 0.000 | -0.358 | 0.721 |
|  |  | **Intercept** | 0.544 | 0.039 | 13.982 | **< 0.001** |
|  | Functional generality | Age | 0.000 | 0.001 | 0.458 | 0.647 |
|  |  | **Bb** | 0.472 | 0.110 | 4.281 | **< 0.001** |
|  |  | Age * Bb | -0.003 | 0.002 | -1.801 | 0.072 |
|  |  | **Spatial autocovariate** | 0.000 | 0.000 | -2.451 | **0.014** |
|  |  | **Intercept** | 0.325 | 0.053 | 6.109 | **< 0.001** |
|  | Taxonomic generality | Age | 0.000 | 0.000 | -0.917 | 0.359 |
|  |  | Bb | -0.024 | 0.045 | -0.533 | 0.594 |
|  |  | Age * Bb | -0.001 | 0.001 | -0.894 | 0.371 |
|  |  | Spatial autocovariate | 0.000 | 0.000 | -0.613 | 0.540 |
|  |  | **Intercept** | 0.266 | 0.023 | 11.780 | **< 0.001** |
| **Colour (fly)** | Species generality | **Age** | -0.002 | 0.001 | -2.848 | **0.004** |
|  |  | **Fy+** | -0.125 | 0.059 | -2.102 | **0.036** |
|  |  | Age * Fy+ | 0.001 | 0.001 | 1.628 | 0.104 |
|  |  | Spatial autocovariate | 0.000 | 0.000 | -0.448 | 0.654 |
|  |  | **Intercept** | 0.521 | 0.045 | 11.527 | **< 0.001** |
|  | Functional generality | Age | 0.001 | 0.001 | 0.646 | 0.518 |
|  |  | **Fy+** | 0.308 | 0.094 | 3.282 | **0.001** |
|  |  | Age * Fy+ | -0.002 | 0.001 | -1.686 | 0.092 |
|  |  | **Spatial autocovariate** | 0.000 | 0.000 | -2.229 | **0.026** |
|  |  | **Intercept** | 0.327 | 0.057 | 5.744 | **< 0.001** |
|  | Taxonomic generality | Age | 0.000 | 0.000 | -0.438 | 0.661 |
|  |  | Fy+ | -0.008 | 0.036 | -0.219 | 0.826 |
|  |  | Age * Fy+ | -0.001 | 0.001 | -1.478 | 0.140 |
|  |  | Spatial autocovariate | 0.000 | 0.000 | -0.547 | 0.584 |
|  |  | **Intercept** | 0.261 | 0.024 | 10.743 | **< 0.001** |
| **Shape** | Conspecific pollen receipt | Age | 0.002 | 0.002 | 0.954 | 0.340 |
|  |  | Mt | -0.265 | 0.162 | -1.636 | 0.102 |
|  |  | **Lt** | -0.443 | 0.166 | -2.665 | **0.008** |
|  |  | **Age * Mt** | 0.006 | 0.003 | 2.225 | **0.026** |
|  |  | Age * Lt | 0.004 | 0.003 | 1.328 | 0.184 |
|  |  | Spatial autocovariate | 0.000 | 0.000 | 0.479 | 0.632 |
|  |  | **Intercept** | -0.221 | 0.004 | -1.787 | **0.074** |
|  | Conspecific pollen receipt | Species Generality (SG) | 0.121 | 0.153 | 0.787 | 0.431 |
|  |  | Functional Generality (FG) | 0.153 | 0.081 | 1.895 | 0.058 |
|  |  | Mt | 0.261 | 0.230 | 1.131 | 0.258 |
|  |  | **Lt** | 0.643 | 0.159 | 4.055 | **< 0.001** |
|  |  | SG * Mt | 0.079 | 0.347 | 0.226 | 0.821 |
|  |  | **SG * Lt** | -0.685 | 0.231 | -2.97 | **0.003** |
|  |  | **FG * Mt** | -0.694 | 0.224 | -3.104 | **0.002** |
|  |  | **FG * Lt** | -5.695 | 0.150 | -3.796 | **< 0.001** |
|  |  | Visiting frequency | -0.022 | 0.018 | -1.203 | 0.229 |
|  |  | **Spatial autocovariate** | 0.000 | 0.000 | 3.391 | **0.001** |
|  |  | Intercept | -0.115 | 0.110 | -1.048 | 0.295 |
| **Colour (bee/fly)** | Conspecific pollen receipt | **Age** | 0.004 | 0.001 | 2.514 | **0.012** |
|  |  | **Bb (Fy+)** | -0.483 | 0.148 | -3.256 | **0.001** |
|  |  | Age * Bb (Fy+) | 0.004 | 0.002 | 1.519 | 0.129 |
|  |  | Spatial autocovariate | 0.000 | 0.000 | 0.098 | 0.922 |
|  |  | **Intercept** | -0.284 | 0.097 | -2.916 | **0.004** |
|  | Conspecific pollen receipt | SG | 0.120 | 0.132 | 0.910 | 0.363 |
|  |  | FG | 0.128 | 0.076 | 1.680 | 0.093 |
|  |  | **Bb (Fy+)** | 0.451 | 0.148 | 3.046 | **0.002** |
|  |  | SG * Bb (Fy+) | -0.340 | 0.212 | -1.607 | 0.108 |
|  |  | **FG * Bb (Fy+)** | -0.570 | 0.134 | -4.257 | **< 0.001** |
|  |  | Visiting frequency | -0.031 | 0.018 | -1.733 | 0.083 |
|  |  | **Spatial autocovariate** | 0.000 | 0.000 | 4.742 | **< 0.001** |
|  |  | Intercept | -0.091 | 0.085 | -1.070 | 0.284 |

**Table S3** Results of generalized linear mixed model analyses for richness, abundance, plant generality for total plant species. Short-tubed, bee-non-blue and fly-y- flowers were used as the baselines for floral shape and bee- and fly- colour groups, respectively. Significant effects are indicated in bold (*P <* 0.05).

| **Response variables** | | **Explanatory variable** | **Coefficients** | | **z** | **P** |
| --- | --- | --- | --- | --- | --- | --- |
|  |  |  | **Estimate** | **SE** |  |  |
| **Shape** | Flower richness | Grassland age (Age) | 0.006 | 0.003 | 1.897 | 0.058 |
|  |  | **Medium-tubed (Mt)** | -0.921 | 0.278 | -3.312 | **0.001** |
|  |  | **Long-tubed (Lt)** | -2.458 | 0.459 | -5.358 | **< 0.001** |
|  |  | Age * Mt | 0.001 | 0.005 | 0.290 | 0.772 |
|  |  | **Age * Lt** | 0.016 | 0.008 | 2.037 | **0.042** |
|  |  | Spatial autocovariate | 0.000 | 0.000 | -0.492 | 0.623 |
|  |  | **Intercept** | 2.686 | 0.154 | 17.480 | **< 0.001** |
|  | Flower abundance | Age | 0.000 | 0.006 | 0.042 | 0.967 |
|  |  | Mt | -0.789 | 0.425 | -1.858 | 0.063 |
|  |  | **Lt** | -2.266 | 0.425 | -5.336 | **< 0.001** |
|  |  | Age * Mt | -0.004 | 0.008 | -0.551 | 0.582 |
|  |  | **Age * Lt** | 0.017 | 0.008 | 2.094 | **0.036** |
|  |  | Spatial autocovariate | 0.000 | 0.000 | 0.688 | 0.492 |
|  |  | **Intercept** | 3.018 | 0.342 | 8.823 | **< 0.001** |
| **Colour (Bee)** | Flower richness | **Age** | 0.007 | 0.003 | 2.496 | **0.013** |
|  |  | **Bee-blue (Bb)** | -1.456 | 0.303 | -4.800 | **< 0.001** |
|  |  | Age * Bb | 0.004 | 0.005 | 0.673 | 0.501 |
|  |  | Spatial autocovariate | 0.000 | 0.000 | -0.470 | 0.638 |
|  |  | **Intercept** | 2.860 | 0.140 | 20.440 | **< 0.001** |
|  | Flower abundance | Age | 0.002 | 0.003 | 0.754 | 0.451 |
|  |  | **Bb** | -2.076 | 0.216 | -9.601 | **< 0.001** |
|  |  | **Age * Bb** | 0.019 | 0.004 | 4.615 | **< 0.001** |
|  |  | Spatial autocovariate | 0.000 | 0.000 | -1.313 | 0.189 |
|  |  | **Intercept** | 3.157 | 0.213 | 14.820 | **< 0.001** |
| **Colour (Fly)** | Flower richness | Age | 0.006 | 0.003 | 1.797 | 0.072 |
|  |  | **Fly-y-plus (Fy+)** | -0.533 | 0.249 | -2.139 | **0.032** |
|  |  | Age * Fy+ | 0.006 | 0.004 | 1.274 | 0.203 |
|  |  | Spatial autocovariate | 0.000 | 0.000 | -0.505 | 0.614 |
|  |  | **Intercept** | 2.597 | 0.161 | 16.124 | **< 0.001** |
|  | Flower abundance | Age | 0.000 | 0.004 | -0.110 | 0.912 |
|  |  | **Fy+** | -1.012 | 0.249 | -4.063 | **< 0.001** |
|  |  | Age * Fy | 0.006 | 0.005 | 1.308 | 0.191 |
|  |  | Spatial autocovariate | 0.000 | 0.000 | 0.344 | 0.731 |
|  |  | **Intercept** | 3.085 | 0.267 | 11.560 | **< 0.001** |
| **Shape** | Species generality | **Age** | -0.001 | 0.000 | -2.329 | **0.020** |
|  |  | Mt | -0.034 | 0.057 | -0.583 | 0.560 |
|  |  | **Lt** | -0.200 | 0.092 | -2.171 | **0.030** |
|  |  | Age * Mt | 0.000 | 0.001 | -0.525 | 0.600 |
|  |  | **Age * Lt** | 0.004 | 0.001 | 3.217 | **0.001** |
|  |  | Spatial autocovariate | 0.000 | 0.000 | -0.380 | 0.704 |
|  |  | **Intercept** | 0.484 | 0.045 | 10.882 | **< 0.001** |
|  | Functional generality | Age | 0.000 | 0.001 | -0.093 | 0.926 |
|  |  | **Mt** | 0.208 | 0.087 | 2.376 | **0.018** |
|  |  | **Lt** | 0.472 | 0.139 | 3.399 | **0.001** |
|  |  | Age * Mt | -0.001 | 0.001 | -0.449 | 0.653 |
|  |  | **Age * Lt** | -0.006 | 0.002 | -3.116 | **0.002** |
|  |  | **Spatial autocovariate** | 0.000 | 0.000 | -2.394 | **0.017** |
|  |  | **Intercept** | 0.389 | 0.048 | 8.110 | **< 0.001** |
|  | Taxonomic generality | Age | 0.000 | 0.000 | -1.162 | 0.245 |
|  |  | Mt | -0.003 | 0.034 | -0.094 | 0.925 |
|  |  | Lt | -0.025 | 0.055 | -0.453 | 0.651 |
|  |  | Age * Mt | 0.000 | 0.001 | 0.409 | 0.682 |
|  |  | Age * Lt | 0.000 | 0.001 | -0.606 | 0.545 |
|  |  | Spatial autocovariate | 0.000 | 0.000 | -0.644 | 0.519 |
|  |  | **Intercept** | 0.259 | 0.021 | 12.064 | **< 0.001** |
| **Colour (Bee)** | Species generality | **Age** | -0.001 | 0.000 | -2.801 | **0.005** |
|  |  | **Bb** | -0.257 | 0.062 | -4.137 | **< 0.001** |
|  |  | **Age * Bb** | 0.002 | 0.001 | 2.334 | **0.020** |
|  |  | Spatial autocovariate | 0.000 | 0.000 | -0.289 | 0.772 |
|  |  | **Intercept** | 0.530 | 0.040 | 13.311 | **< 0.001** |
|  | Functional generality | Age | 0.000 | 0.001 | -0.462 | 0.644 |
|  |  | **Bb** | 0.446 | 0.094 | 4.743 | **< 0.001** |
|  |  | **Age * Bb** | -0.003 | 0.001 | -2.273 | **0.023** |
|  |  | **Spatial autocovariate** | 0.000 | 0.000 | -2.299 | **0.021** |
|  |  | **Intercept** | 0.380 | 0.045 | 8.469 | **< 0.001** |
|  | Taxonomic generality | Age | 0.000 | 0.000 | -0.935 | 0.350 |
|  |  | Bb | -0.032 | 0.039 | -0.825 | 0.410 |
|  |  | Age * Bb | 0.000 | 0.001 | -0.519 | 0.604 |
|  |  | Spatial autocovariate | 0.000 | 0.000 | -0.710 | 0.478 |
|  |  | **Intercept** | 0.264 | 0.020 | 13.234 | **< 0.001** |
| **Colour (Fly)** | Species generality | **Age** | -0.001 | 0.000 | -2.750 | **0.006** |
|  |  | **Fy+** | -0.112 | 0.052 | -2.151 | **0.031** |
|  |  | Age * Fy+ | 0.001 | 0.001 | 1.704 | 0.088 |
|  |  | Spatial autocovariate | 0.000 | 0.000 | -0.349 | 0.727 |
|  |  | **Intercept** | 0.509 | 0.044 | 11.538 | **< 0.001** |
|  | Functional generality | Age | 0.000 | 0.001 | -0.357 | 0.721 |
|  |  | **Fy+** | 0.232 | 0.079 | 2.945 | **0.003** |
|  |  | Age * Fy+ | -0.002 | 0.001 | -1.507 | 0.132 |
|  |  | **Spatial autocovariate** | 0.000 | 0.000 | -2.101 | **0.036** |
|  |  | **Intercept** | 0.395 | 0.049 | 8.112 | **< 0.001** |
|  | Taxonomic generality | Age | 0.000 | 0.000 | -0.706 | 0.480 |
|  |  | Fy+ | -0.017 | 0.030 | -0.557 | 0.577 |
|  |  | Age * Fy+ | 0.000 | 0.000 | -0.660 | 0.509 |
|  |  | Spatial autocovariate | 0.000 | 0.000 | -0.729 | 0.466 |
|  |  | **Intercept** | 0.259 | 0.022 | 11.983 | **< 0.001** |

**Table S4** Correlations between flower and pollinator composition. Significant effects are indicated in bold (*P < 0.05*).

| **Native** | Bee colour group | | Fly colour group | |
| --- | --- | --- | --- | --- |
|  | Flower PC1 | Flower PC2 | Flower PC1 | Flower PC2 |
| Pollinator PC1 | **0.710** | 0.186 | **0.644** | 0.426 |
| Pollinator PC2 | -0.121 | 0.266 | -0.138 | 0.044 |
| **Total** |  |  |  |  |
| Pollinator PC1 | **0.727** | 0.124 | **0.687** | -0.162 |
| Pollinator PC2 | -0.063 | 0.188 | 0.004 | 0.029 |
